# Abdominal-B neurons selectively drive vibrations in *Drosophila*

**DOI:** 10.64898/2026.07.12.737445

**Authors:** Elsa Steinfath, Kimia Alizadeh, Jan Clemens

## Abstract

Male *Drosophila* courtship includes two communication signals: airborne song and substrate-borne vibrations. While the neural control of song has been extensively characterized, little is known about the circuits underlying vibration production. Here, we identify neurons expressing the Hox gene abdominal-B (*abdB*) as a driver of vibration production. Optogenetic activation of *abdB* neurons selectively elicited vibrations in both males and females without inducing courtship song, whereas silencing these neurons did not impair vibration production during natural courtship. The vibration-driving *abdB* neurons are neither *doublesex*-nor *fruitless*-positive, defining a previously unrecognized component of the courtship circuit. Although *abdB* activation produced only stimulus-locked vibrations, co-activation of the persistence-promoting neuron cluster pCd converted this transient signal output into sustained vibration trains. Together, our results identify a dedicated pathway for vibration production and show that signal identity and persistence can be independently specified by distinct circuit components.

## Introduction

Social behavior emerges from neural computations distributed across multiple levels of the nervous system. Sensory neurons detect external cues and self-motion [1–4], ascending neurons and interneurons relay this information to the brain [1, 5–7], where central circuits integrate sensory information with internal state to select appropriate actions [8–13]. Descending neurons then transmit motor commands to the ventral nerve cord, where pattern-generating circuits and motor neurons produce behavior [14–19]. Understanding how distinct motor programs are selected and coordinated across this hierarchy remains a central goal in systems neuroscience.

*Drosophila* courtship provides an excellent model for studying this problem. During courtship, males continuously integrate visual, olfactory, gustatory and mechanosensory information from the female with information about their own movement state [1, 7, 20–22]. These signals are processed by dedicated courtship circuits that interact with internal-state networks and descending pathways to generate a coordinated sequence of behaviors including pursuit, tapping, courtship signal production and copulation attempts [1, 2, 9, 12, 23–28].

A central feature of courtship is the production of multiple communication signals. Most work has focused on courtship song because its production is dynamically regulated by social context [29–31] and its underlying circuitry has been extensively characterized [15, 27]. However, males also produce substrate-borne vibrations [32, 33]. Whereas song is generated by unilateral wing vibration and consists of pulse and sine song [34], vibrations are produced without wing movement, are associated with abdominal quivering, and occur in trains with inter-vibration intervals of approximately 150–200 ms [33, 35]. Both signals are detected by females and influence mating decisions [22, 36, 37], suggesting that successful courtship requires the coordinated control of distinct signaling programs in the male.

In contrast to the extensive understanding of song production, little is known about the neural control of vibration. Vibration production has been broadly linked to neurons expressing the sex-determination genes *doublesex* (*dsx*) and *fruitless* (*fru*) [33]. Recent work showed that the central brain neurons P1a and pC2l coordinate song and vibration production and that persistent vibration can be explained by recurrent circuit dynamics [35]. These findings established central mechanisms that regulate vibration, but left two key questions unresolved: which neurons outside the central brain drive vibration, and how persistent vibrational output is generated.

Here we identify neurons expressing the Hox gene *abdominal-B* (*abdB*) [38] as a previously unknown component of the vibration circuit. We show that activation of *abdB* neurons, whose somata reside primarily in the abdominal ganglion of the ventral nerve cord [37], is sufficient to evoke vibrations in both males and females but is not required for vibration production during natural courtship. Furthermore, we show that the vibration-driving *abdB* neurons are neither *dsx*-nor *fru*-positive, and that co-activation with the persistence-promoting neuron cluster pCd converts stimulus-locked vibrations into sustained vibration trains. Together, these findings identify a dedicated circuit for vibration production and show how its output is shaped by central circuits controlling courtship persistence.

## Results

### Activation of *abdB* neurons selectively drives vibrations

To identify neurons outside the central brain that can drive vibration production, we optogenetically activated several candidate neuronal populations, including *dsx, fru* and *abdB*, in solitary males while recording song and vibration using our previously established multi-microphone assay [35], modified from [29, 39]. Flies walked on a thin paper substrate covering the microphones, allowing simultaneous detection of courtship song and vibrations.

We focused on *abdB*, which labels approximately 280 neurons with somata predominantly located in the abdominal ganglion of the ventral nerve cord and has previously been implicated in female receptivity [37] (Fig. 1A). Optogenetic activation of *abdB* reliably elicited vibrations in solitary males (Fig. 1B). Vibration trains began shortly after stimulus onset, gradually increased in probability, persisted throughout stimulation, and terminated with the stimulus. This stimulus-locked behavior was observed across stimulation durations of 5, 10 and 20 s (Fig. 1C–E).

**Figure 1:**
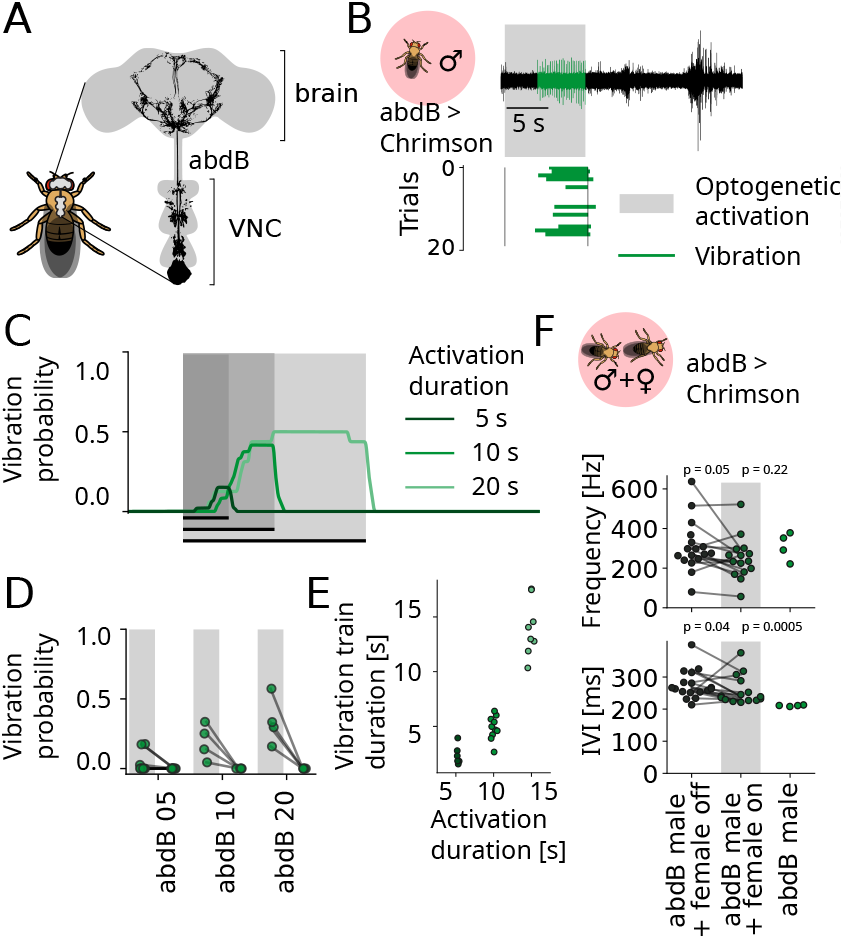
*abdB* neurons drive vibration in males. **A** Schematic of *abdB* expression (from [37]). Most neuronal somata are located in the abdominal ganglion of the ventral nerve cord (VNC), and only a small number of neurons project to the central brain. **B** Example microphone recording (top) and trial raster (bottom) showing vibrations (green) elicited by optogenetic activation of *abdB* neurons in solitary males. **C** Mean vibration probability during optogenetic activation of *abdB* neurons in solitary males using 5, 10 and 20 s stimulation (14 mW/cm^2^; N=4 flies, 4–6 trials per fly). **D** Mean vibration probability during (grey background) and outside (white) optogenetic stimulation for each stimulation duration (same data as C). **E** Duration of individual vibration trains elicited by optogenetic activation for each stimulation duration (same data as C and D). **F** Carrier frequency (top) and inter-vibration interval (IVI, bottom) of vibrations produced by courting *abdB*>Chrimson males (14 and 29 mW/cm^2^; N=17 couples, 8 trials per couple) during and outside optogenetic stimulation, compared with vibrations produced by solitary *abdB*>Chrimson males (same flies as C–E). Data from courting males failed the Shapiro–Wilk test for normality. Paired comparisons between LED on and LED off were performed using a Wilcoxon signed-rank test. Comparisons between solitary males and courting males during optogenetic stimulation were performed using a Mann–Whitney U test. Statistical significance was assessed using a Bonferroni-corrected significance threshold of *P <* 0.0125.

The optogenetically evoked vibrations closely resembled naturally produced vibrations (Fig. 1F). Their carrier frequency was indistinguishable from those recorded during courtship, whereas inter-vibration intervals were slightly shorter than those of courting *abdB* males but remained within the range previously reported for wild-type *D. melanogaster* males [33, 35].

Unlike the central brain neuron clusters P1a and pC2l, whose activation induces both song and vibrations [35], activation of *abdB* never elicited courtship song. To our knowledge, *abdB* is therefore the first driver line reported to selectively evoke vibrations without co-activating song.

### *abdB* is sufficient but not required for vibration production

We next asked whether *abdB* neurons are required for vibration production during natural courtship. To this end, we optogenetically silenced *abdB* neurons using the light-gated chloride channel GtACR1 [40] in males courting wild-type females. Inactivation of *abdB* neurons had no detectable effect on vibration production (Fig. 2A). Males were equally likely to produce vibrations during and outside photoinhibition (Fig. 2B–C), and vibration carrier frequency and inter-vibration interval remained unchanged (Fig. 2D). Thus, although activation of *abdB* neurons is sufficient to evoke vibrations, these neurons are not required for vibration production during normal courtship.

**Figure 2:**
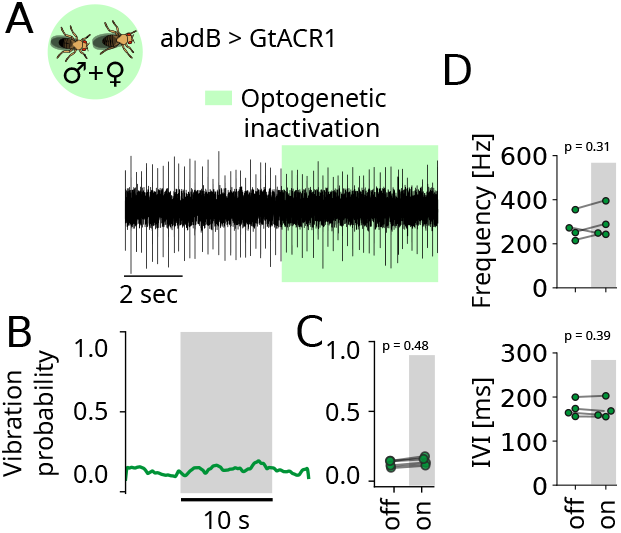
*abdB* neurons are not required for male vibration. **A** Example microphone recording showing vibrations produced by a male courting a wild-type female around the onset of optogenetic inactivation of *abdB* neurons (green). **B** Mean vibration probability during optogenetic inactivation of *abdB* neurons in males courting wild-type females (6 mW/cm^2^; N=4 fly couples, 30 trials per couple). **C** Mean vibration probability during and outside optogenetic inactivation (same data as B). **D** Carrier frequency (top) and inter-vibration interval (IVI, bottom) of vibrations produced during and outside optogenetic inactivation (same data as B and C). Data passed the Shapiro–Wilk test for normality. Paired comparisons between LED on and LED off were performed using paired *t*-tests. Statistical significance was assessed using a Bonferroni-corrected significance threshold of *P <* 0.017.

### *abdB* activation reveals a latent vibration circuit in females

To further localize the neurons responsible for the vibration phenotype, we restricted *abdB* expression using *teashirt* (tsh)-GAL80, which reduces the labeled population by approximately half [37]. Optogenetic activation of this *abdB∩tsh* population continued to elicit stimulus-locked vibrations in solitary males with dynamics similar to those observed using the full *abdB* driver (Fig. 3A–B), indicating that the responsible neurons are contained within this subset.

**Figure 3:**
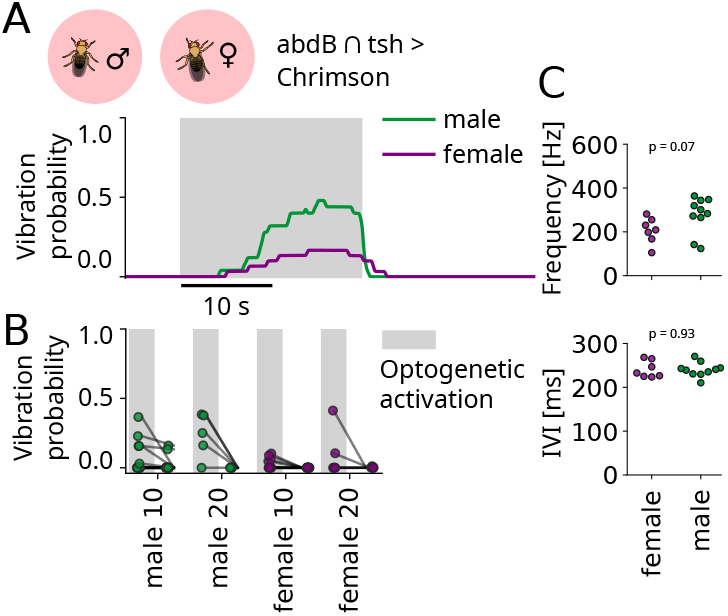
*abdB* neurons drive vibration in females. **A** Mean vibration probability during 20 s optogenetic activation of *abdB∩tsh* neurons in solitary males (green; N=5 flies) and females (purple; N=6 flies) (14 mW/cm^2^; 5–6 trials per fly). **B** Mean vibration probability during and outside optogenetic activation in solitary males (green; N=5 flies) and females (purple; N=6 flies) for 10 and 20 s stimulation (14 mW/cm^2^; 5–6 trials per fly). **C** Carrier frequency (top) and inter-vibration interval (IVI, bottom) of vibrations produced by solitary *abdB∩tsh*>Chrimson males and females during optogenetic activation. Data were pooled across stimulation durations and intensities (males: 14 mW/cm^2^; females: 14 or 29 mW/cm^2^). Data passed the Shapiro–Wilk test for normality. Male and female data were compared using independent *t*-tests. Statistical significance was assessed using a Bonferroni-corrected significance threshold of *P <* 0.025.

Substrate-borne vibrations have so far only been described in male flies [33, 41, 42]. Surprisingly, activation of the same *abdB∩tsh* population also elicited vibrations in solitary females (Fig. 3A–B). Although females responded less reliably than males (7/12 females versus 10/11 males), the vibrations they produced closely resembled those of males in carrier frequency and inter-vibration interval (Fig. 3C). Thus, the neuronal circuitry capable of generating vibrations is present in both sexes and can be recruited by activation of *abdB* neurons.

### The vibration-driving *abdB* neurons are neither *dsx*-nor *fru*-positive

Because courtship behaviors are largely controlled by neurons expressing the sex-determination genes *doublesex* (*dsx*) and *fruitless* (*fru*) [43], we asked whether the *abdB* neurons driving vibrations belong to either of these populations. This appeared plausible because vibration production has previously been linked to *dsx* and *fru* neurons [33], and the ascending neuron vAB3 co-expresses *abdB* and *fru* and projects to the central courtship circuit [1, 44, 45].

As expected, optogenetic activation of either *dsx* or *fru* elicited vibrations in solitary males. However, the resulting behavioral dynamics differed markedly from those evoked by *abdB*. Whereas *abdB* activation generated stimulus-locked vibrations, activation of *dsx* produced long-lasting vibration trains that started after stimulus offset and persisted (Fig. 4A). Activation of *fru* evoked only sparse vibrations, again after stimulation (Fig. 4A). Both driver lines also induced courtship song [33, 46, 47], in contrast to the selective vibration phenotype observed following *abdB* activation.

**Figure 4:**
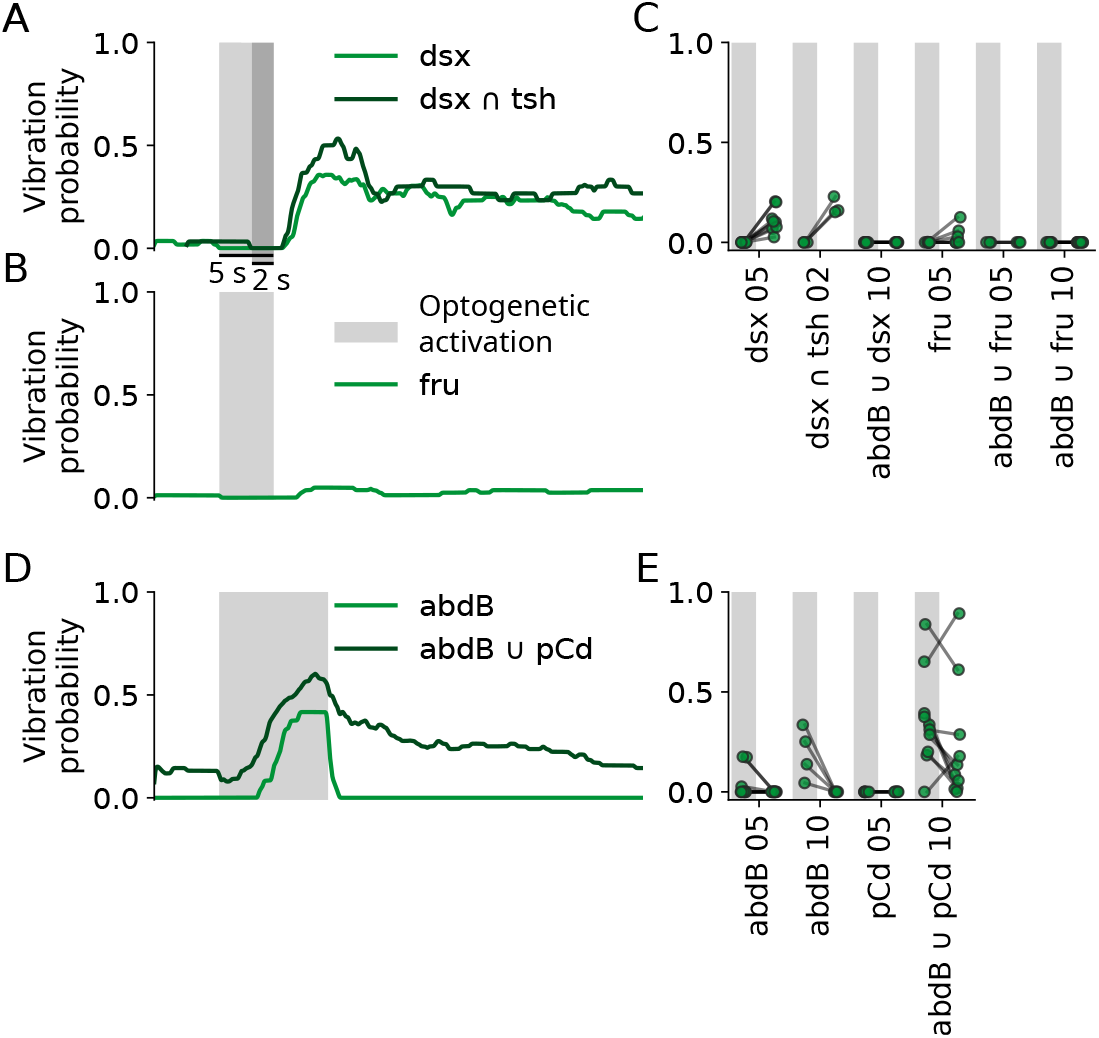
Sex-specific neuron populations modulate vibration persistence in males. **A** Mean vibration probability during optogenetic activation of *dsx* (5 s stimulation, 19 mW/cm^2^; N=8 flies, 7 trials per fly) and *dsx∩tsh* (2 s stimulation, 14 mW/cm^2^; N=3 flies, 10 trials per fly) in solitary males. **B** Mean vibration probability during optogenetic activation of *fru* neurons in solitary males (14 mW/cm^2^; N=7 flies, 6–12 trials per fly). **C** Mean vibration probability during and outside optogenetic activation of *dsx* (N=8 flies), *dsx∩tsh* (N=3 flies), *abdB∪dsx* (N=7 flies), *fru* (N=7 flies), and *abdB∪fru* (N=10 flies). **D** Mean vibration probability during optogenetic activation of *abdB* neurons (14 mW/cm^2^; N=4 flies, 4–6 trials per fly) or co-activation of *abdB* and *pCd* neurons (14 mW/cm^2^; N=10 flies, 6–8 trials per fly) in solitary males. **E** Mean vibration probability during and outside optogenetic activation of *abdB* (N=4 flies), *pCd* (N=4 flies), and *abdB*+*pCd* co-activation (N=10 flies).

To determine whether the vibration-driving *abdB* neurons are part of either population, we activated the genetic intersections *abdB∩dsx* and *abdB∩fru* (Fig. 4C). Neither intersection elicited vibrations, indicating that the neurons responsible for the *abdB* phenotype are neither *dsx*-nor *fru*-positive. Together, these results identify the vibration-driving *abdB* neurons as a genetically distinct population from the canonical sex-specific courtship circuitry.

### Vibration production elicited by *abdB* requires the central brain

The *abdB* driver labels neurons whose somata are almost exclusively located in the abdominal ganglion of the ventral nerve cord [37]. We therefore asked whether activation of these neurons drives vibrations locally within the ventral nerve cord or requires signaling through the brain.

To distinguish between these possibilities, we optogenetically activated *abdB* neurons in decapitated males. In contrast to intact flies, decapitated males never produced vibrations (0/5 decapitated males vs. 10/11 intact males, Fig. 1), indicating that the vibration phenotype requires an intact brain.

As a positive control, we activated *dsx* neurons in decapitated males. Consistent with previous reports [47], activation still elicited wing extension and, in several animals, courtship song. However, none of the flies produced vibrations (N=6, data not shown). Thus, although activation of *dsx* neurons can directly engage components of the song motor program in the absence of the brain, neither *dsx* nor *abdB* activation is sufficient to drive vibrations without central brain circuits.

These findings indicate that *abdB*-evoked vibration production is gated by the brain, consistent with a role for ascending neurons that relay activity from the abdominal ganglion to central courtship circuits before descending pathways initiate the motor program.

### pCd converts transient *abdB*-evoked vibrations into persistent signal output

Activation of *abdB* elicited vibrations only during optogenetic stimulation and never induced courtship song. In contrast, all previously identified drivers of vibration production, including P1a and pC2l, evoke vibrations that outlast stimulation [35]. We therefore asked whether persistence can be added to the *abdB*-evoked signaling program by co-activating neurons involved in courtship persistence.

The *dsx*-positive neuron cluster pCd forms part of a recurrent circuit that maintains courtship motivation and persistent social behaviors, including persistent singing [12, 25, 48]. As expected, optogenetic activation of pCd alone did not elicit song or vibrations in solitary males (Fig. 4E, data for song not shown). Remarkably, however, simultaneous activation of *abdB* and pCd converted the transient *abdB* phenotype into long-lasting vibration trains that persisted well beyond stimulus offset (Fig. 4D, E).

Unlike activation of P1a or pC2l, co-activation of *abdB* and pCd remained selective for vibrations and did not evoke courtship song (data not shown). Thus, pCd does not specify the signaling program itself but prolongs signal output generated by *abdB*. These results demonstrate that vibration identity and persistence are controlled by separable circuit components: *abdB* provides a vibration-specific signaling drive, whereas pCd endows this signal output with persistence.

## Discussion

Courtship requires males to select, coordinate and sustain multiple courtship signals. While the neural control of courtship song is increasingly well understood, much less is known about the circuitry underlying substrate-borne vibrations. Our results suggest that vibration production is organized hierarchically. We identify an *abdB*-positive pathway that selectively accesses the vibration signaling program, reveal that this program can also be recruited in females, and show that central courtship-state circuits regulate whether vibration output is transient or persistent.

### A vibration-specific access pathway

The observation that *abdB* neurons are sufficient (Fig. 1), but not required (Fig. 2), for vibration production suggests that they are not obligatory components of the vibration pathway. Rather, *abdB* appears to provide one of several routes by which downstream vibration-generating circuitry can be engaged. Unlike previously described vibration-driving populations, activation of *abdB* selectively evokes vibrations without courtship song and produces stimulus-locked rather than persistent behavioral output (Fig. 1). We therefore propose that *abdB* provides relatively selective access to the vibration program, whereas the broader behavioral state associated with courtship is supplied by central circuits.

Unexpectedly, the vibration-driving *abdB* neurons are genetically distinct from the canonical *fru*- and *dsx*-positive courtship populations (Fig. 4A–C). Together with the finding that activation of the same pathway elicits vibrations in females (Fig. 3), this suggests that the motor substrate underlying vibration is not itself strongly sex-specific. Instead, sex specificity may arise primarily from how this motor program is recruited during courtship, despite the extensive sexual dimorphism of the abdominal ganglion and central courtship circuits [49, 50].

This idea is consistent with existing work showing that many male courtship behaviors rely on latent circuitry that is also present in females. Manipulating the sex-determination pathway through *fruitless* or *doublesex*, or directly activating central courtship neurons, can elicit male-typical behaviors in females, including chasing, wing extension and even courtship song [43, 51–53]. Our findings extend this principle to substrate-borne vibrations. Unlike these previous studies, however, female vibration is recruited through an *abdB*-positive population associated with the abdominal ganglion rather than by manipulating central courtship circuitry, suggesting that the downstream motor machinery is shared between the sexes.

### *abdB*-evoked vibrations require central brain circuits

Although *abdB* neurons reside primarily in the abdominal ganglion, their activation requires an intact brain to elicit vibrations. This finding argues against a purely local abdominal or VNC mechanism and instead suggests that vibration production depends on communication between the abdominal ganglion and central courtship circuits. Importantly, however, the decapitation experiment alone does not establish the role of the brain in this process. At one extreme, descending pathways could simply provide a permissive courtship-state signal that enables VNC circuits to respond to *abdB*-driven input, without actively shaping the motor program itself.

However, the pCd co-activation experiments suggest that the contribution of central brain circuits extends beyond such permissive gating (Fig. 4D–E). Previous work established pCd as a central component of the persistent courtship-state network downstream of P1, where it maintains courtship motivation and prolongs behaviors such as wing extension and likely also singing [12, 24, 25, 48]. We extend this model by showing that pCd can acutely convert transient *abdB*-evoked vibrations into persistent behavioral output without altering their identity. Because pCd alone does not evoke vibrations, it appears to regulate the persistence, rather than the identity, of an already active signaling program. Thus, central brain circuits not only permit the expression of vibration but also actively regulate its temporal dynamics.

These findings support a hierarchical organization of the vibration circuit in which an *abdB*-dependent pathway provides selective access to the vibration signaling program, while central brain circuits determine whether, and for how long, that program is expressed. Whether the relevant *abdB* neurons are ascending neurons that recruit these central circuits, or local VNC neurons whose output depends on descending signals, remains unresolved. The failure of the *abdB∩fru* intersection to evoke vibrations argues against the known ascending neuron vAB3 as the principal pathway linking the abdominal ganglion to the central courtship network [1], pointing instead to previously unidentified *abdB*-positive neurons.

### Open questions

A major challenge for future work will be to identify the vibration-driving neurons within the *abdB* population. The current driver labels approximately 140 neurons after *tsh* subtraction [37], likely including a mixture of local interneurons, ascending neurons and motor-related neurons. Inter-sectional genetic strategies or stochastic sparse-labeling approaches such as SPARC [54] should enable identification of the minimal population sufficient to drive vibration production.

A second question is how *abdB* activity engages central courtship circuits. If the relevant neurons are ascending neurons, they may relay sensory or behavioral-state information from the abdominal ganglion to the brain. Alternatively, they may form part of local VNC circuitry whose output depends on descending courtship-state signals. Functional imaging combined with connectome-guided circuit reconstruction should distinguish between these possibilities and reveal how abdominal signals interact with central circuits controlling courtship state and behavioral persistence.

A third question concerns the relationship between the vibration signaling program and the recently described nested architecture of the song central pattern generator [27]. The ability of *abdB* activation to evoke vibrations in the complete absence of song suggests that vibration production is not simply another output mode of the song circuit. Instead, song and vibrations are likely generated by distinct but interacting motor circuits that converge on shared motor neurons. Identifying where these circuits interact will be important for understanding how flies rapidly switch between and coordinate multiple communication signals during courtship.

Finally, the downstream circuitry that generates vibrations remains unknown. Recent work has begun to elucidate the biomechanics of vibration production, implicating rhythmic abdominal tremulation and longitudinal stretch receptors in this behavior [55]. However, it remains unclear whether abdominal movements are themselves required for vibration generation or instead reflect a correlated motor output. Identifying the downstream targets of the *abdB* neurons will provide an entry point for dissecting how descending commands are transformed into the rhythmic motor activity underlying substrate-borne vibrations.

## Conclusion

Our findings support a modular organization of vibration control, in which distinct circuit modules govern signal-program identity, behavioral state and behavioral persistence. Identifying the neurons that link these modules will provide an opportunity to understand how central courtship circuits coordinate multiple communication signals and, more broadly, how distributed neural circuits assemble complex social behaviors from reusable motor components.

## Acknowledgments

We thank Frank Kötting, Stephan Löwe from the ENI workshop for help with designing behavioral chambers, Gesa Hoffmann, Jan Schöning, Christiane Becker, Rüdiger Ludwig and Matthias Weyl for technical and adminstrative assistance. We thank Peter Andolfatto, Andre Fiala, Martin Göpfert, Mala Murthy, Janelia flylight and Bloomington stock center for gifts of flies. This work was funded via an Emmy Noether Grant (Project number 329518246) and an ERC Starting Grant (Grant agreement No. 851210) to JC.

## Author contributions

- Conceptualization - ES, JC
- Animals and behavioral experiments - ES, KA
- Analysis - ES
- First draft - ES, JC

## Methods

### Fly strains and rearing

Flies were kept on a 12:12 hour dark:light cycle, at 25°C and 60% humidity. Flies were collected as virgins within 8 hours after eclosion, separated by sex, and then housed in groups of 3-15 flies.

### Behavioral setups

The behavioral chamber measured 44 mm in diameter and 1.9 mm in height; chamber and lid were machined from transparent acrylic. Chamber lids were coated with Sigmacote (Sigma-Aldrich) to prevent flies from walking on the ceiling, and kept under a fume hood to dry for at least 10 minutes.

The floor of the chamber was tiled with 16 microphones (Knowles NR-23158) that were embedded into a custom-made PCB board (design modified from Coen et al. [29]). The microphones were covered with a thin, white paper for the flies to walk on and to record sound and vibration. Microphone signals were amplified using a custom-build amplifier [39] and digitized using a data acquisition card (National Instruments Pcie-6343) at sampling rate of 10 kHz.

Fly behavior was recorded using a USB camera (FLIR flea3 FL3-U3-13Y3M-C, 100 frames per second (fps), 912 × 920 pixels), equipped with an 35 mm f1.4 objective (Thorlabs MVL35M1). The chamber was illuminated with weak blue light (470 nm). A 500 nm shortpass filter (Edmund Optics, 500 nm 50 mm diameter, OD 4.0 Shortpass Filter) filtered out green (525 nm) and red (625 nm) wavelengths used for optogenetic experiments.

Synchronous recordings of audio, video, and delivery of optogenetic stimuli was controlled using custom software https://janclemenslab.org/etho.

### Behavioral assays

For all experiments, 3 to 7 day old naive males and virgin females were used. Flies were introduced gently into the chamber using an aspirator. All recordings were performed during the flies’ morning activity peak and started within 120 minutes of the incubator lights switching on.

In experiments using decapitated males, flies were cold-anesthetized and the head was cut using a sharp razor blade directly before the experiment.

### Optogenetics

Flies were kept for 3-7 days prior to the experiment on food containing retinal (1 ml all-trans retinal (Sigma-Aldrich) solution (100 mM in 95% ethanol) per 100 ml food). To prevent the degradation of the retinal and continuous neural activation, the vials were wrapped in aluminium foil.

For neural inactivation, we used the GtACR1 channel [40, 58], which was excited using a green LED (625 nm). For inactivation of abdB (Fig. 3D-G) we used an LED intensity of 14 mW/cm^2^. Experiment consisted of 30 trials of optogenetic stimulation. Each trial started with 10 s stimulation (green LED on) followed a pause of 20 s. For neural activation, we used the CsChrimson channel [56], which was activated using a red LED (625 nm). Activation was performed with LED intensities, stimulus duration, trial length and number indicated in Tables 3-2.

**Table 1:** Fly lines used.

| Figs | Name | Genotype | Source | Provided by |
| --- | --- | --- | --- | --- |
| 1, 2 | wild-type | *Drosophila melanogaster* NM91 | [29] | Peter Andolfatto |
| 1-4 | abdB | y[1] w[*]; Pw[+mW.hs]=GawBAbd-B[LDN]-GAL4/TM6B, Tb[1] | [37] | BL 55848 |
| 1, 3, 4 | CsChrimson | UAS20x-CsChrimson.mVenus (II) | [56] | CsChrimson by André Fiala |
| 2 | GtACR1 | UAS-GtACR1.d.EYFP(attP2) | [40] |  |
| 3 | abdB ∩ tsh | ; tsh-GAL80/UAS20x-CsChrimson.mVenus (II); Pw[+mW.hs]=GawBAbd-B[LDN]-GAL4/+ | [37, 57] | tsh-GAL80 by Martin Göpfert |
| 4 | abdB ∪ dsx | y[1] w[*]; Pw[+mW.hs]=GawBAbd-B[LDN]/TM6B, Tb[1] x ; UAS<stop>CsChrimson-mVenus/CyO; dsx-FLP/TM6B | [37] | dsx FLP by Mala Murthy |
| 4 | abdB ∪ fru | y[1] w[*]; Pw[+mW.hs]=GawBAbd-B[LDN]/TM6B, Tb[1] x ; UAS<stop>CsChrimson-mVenus/CyO; fru-FLP/TM6B | [37] | fru FLP by Mala Murthy |
| 4 | dsx | ; +/s; dsx-GAL4/20XUAS-CsChrimson.mVenus |  |  |
| 4 | fru | ; 20XUAS-CsChrimson.mVenus (II)/+; fru-GAL4/+ |  |  |
| 4 | dsx ∩ tsh | ; tsh-GAL80/UAS20x-CsChrimson.mVenus (II); dsx-GAL4/sb | [27, 57] | |
| 4 | pCd | ; UAS20x-CsChrimson.mVenus (II)/+; PGMR41A01-GAL4attP2/+ | [25] | BL 39425 |
| 4 | abdB + pCd | ; UAS20x-CsChrimson.mVenus (II)/+; PGMR41A01-GAL4attP2/Pw[+mW.hs]=GawBAbd-B[LDN]-GAL4 | [25] | BL 39425 |

**Table 2:**
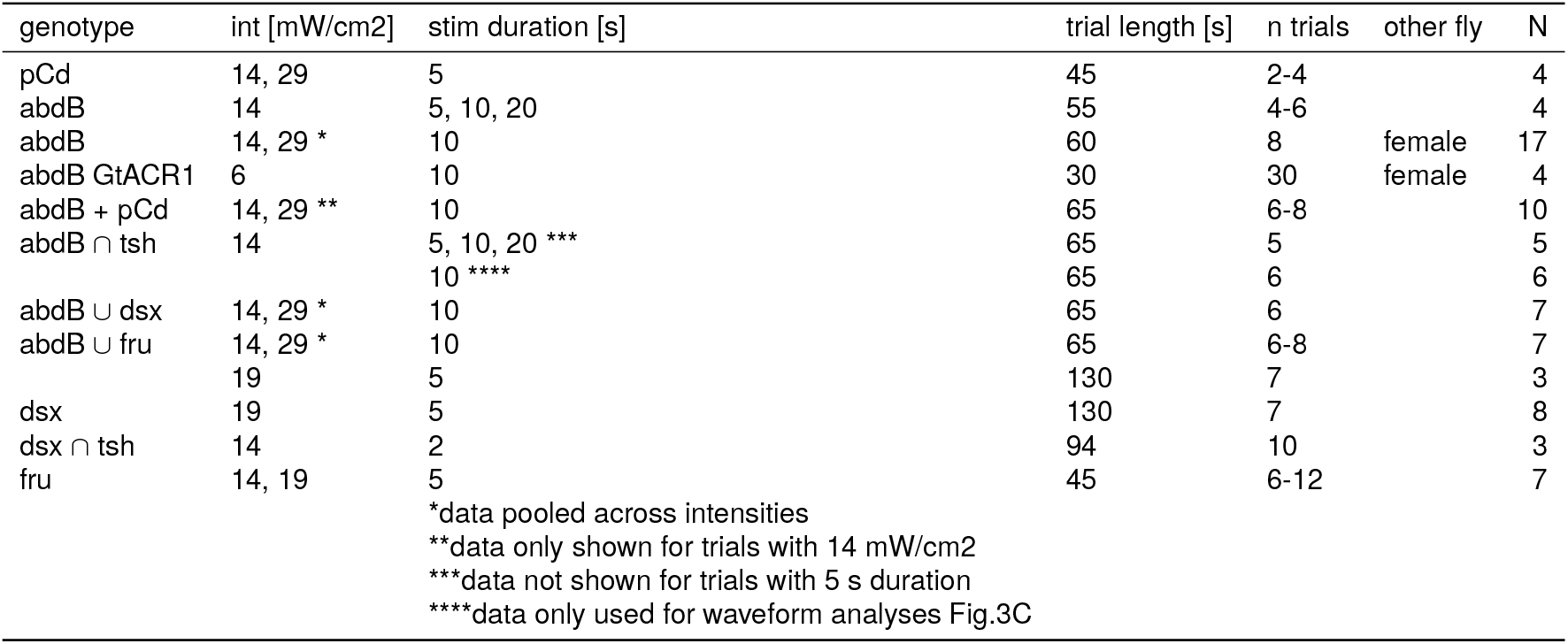
Optogenetic activation regimes for male flies.

**Table 3:**
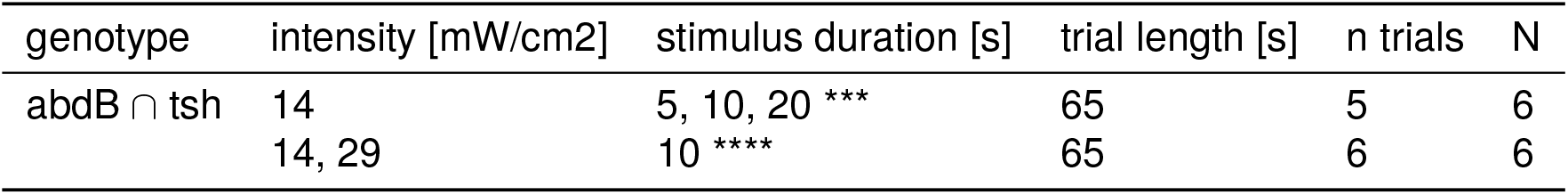
Optogenetic activation regimes for female flies.

**Table 4:**
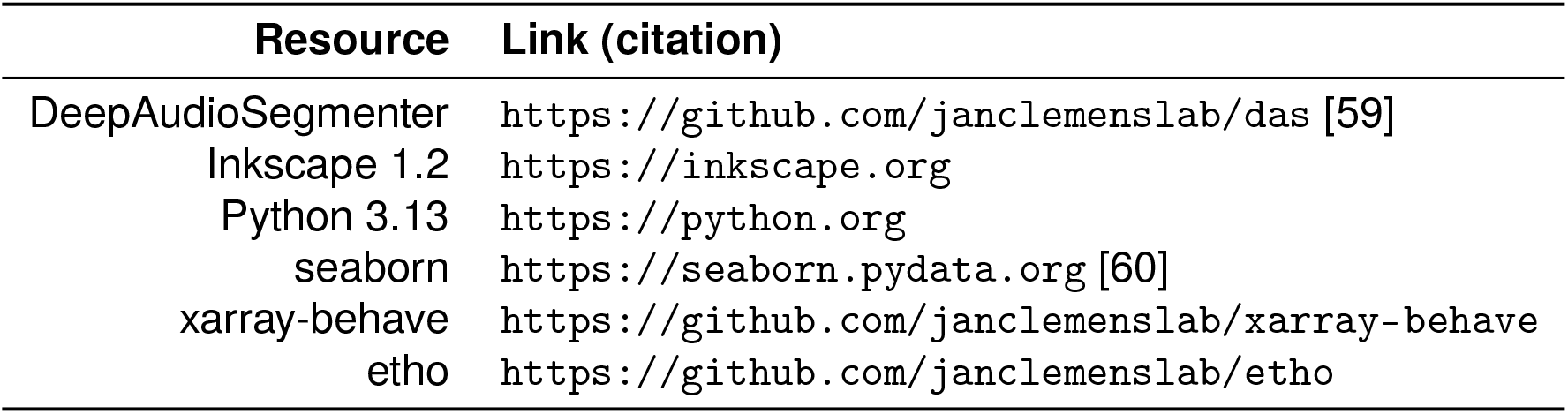
Open source software used.

### Analysis of microphone signals

Vibration signals were manually annotated using the graphical user interface of DAS [59]. For optogenetic manipulation of abdB inactivation (Fig. 3D-G) and abdB activation in males courting a female (Fig. 1F), the annotators were blind to experimental condition.

*Vibration trains* were defined as groups of pulses with an interval less than 400 ms (2–2.5 times of the modal interval).

*Vibration probabilities* for experiments with optogenetic neural activation or inactivation, are given as the fraction of trials during which vibration trains were produced, pooled data across flies. All traces shown for optogenetic experiments are smoothed with a Gaussian window with a standard deviation of 0.1 s. For statistics and scatter plots we computed the mean across trials pooled for each fly.

#### Waveform analysis

Microphone recordings of the same signal on different microphones were regarded as one and the waveform was taken from the loudest channel. To obtain vibration pulses, we extracted 60 ms long waveforms from all time points annotated as vibration signal. To remove variability in the waveforms arising from the position and distance of the singing male from the microphone, we normalized waveforms as described before [34]: we divided the raw waveforms by their norm, centered them to their peak energy and flipped their sign such that the average of the 1 ms preceding the pulse center was positive. Frequencies were obtained by taking the center of mass from the Fourier transformed waveforms. Inter-vibration intervals (IVIs) were calculated as the time differences between consecutive annotated vibration signals. To exclude intervals spanning adjacent vibration trains, intervals exceeding twice the median IVI across all experiments were excluded from the analysis.

